# DNA duplication-mediated activation of a two-component regulatory system serves as a bet-hedging strategy for *Burkholderia thailandensis*

**DOI:** 10.1101/2024.12.09.627470

**Authors:** Lillian C. Lowrey, Katlyn B. Mote, Peggy A. Cotter

**Affiliations:** Department of Microbiology and Immunology, The University of North Carolina at Chapel Hill, School of Medicine, Chapel Hill, North Carolina, USA

## Abstract

*Burkholderia thailandensis* strain E264 (*Bt*E264) and close relatives stochastically duplicate a 208.6 kb region of chromosome I via RecA-dependent recombination between two nearly identical insertion sequence elements. Because homologous recombination occurs at a constant, low level, populations of *Bt*E264 are always heterogeneous, but cells containing two or more copies of the region (Dup+) have an advantage, and hence predominate, during biofilm growth, while those with a single copy (Dup–) are favored during planktonic growth. Moreover, only Dup+ bacteria form ‘efficient’ biofilms within 24 hours in liquid medium. We determined that duplicate copies of a subregion containing genes encoding an archaic chaperone-usher pilus (*aplFABCDE*) and a two-component regulatory system (*bubSR*) are necessary and sufficient for generating efficient biofilms and for conferring a selective advantage during biofilm growth. BubSR functionality is required, as deletion of either *bubS* or *bubR*, or a mutation predicted to abrogate phosphorylation of BubR, abrogates biofilm formation. However, duplicate copies of the *aplFABCDE* genes are not required. Instead, we found that BubSR controls expression of *aplFABCDE* and *bubSR* by activating a promoter upstream of *aplF* during biofilm growth or when the 208.6 kb region, or just *bubSR*, are duplicated. Single cell analyses showed that duplication of the 208.6 kb region is sufficient to activate BubSR in 75% of bacteria during planktonic (BubSR ‘OFF’) growth conditions. Together, our data indicate that the combination of deterministic two-component signal transduction and stochastic, duplication-mediated activation of that TCS form a bet-hedging strategy that allows *Bt*E264 to survive when conditions shift rapidly from those favoring planktonic growth to those requiring biofilm formation, such as may be encountered in the soils of Southeast Asia and Northern Australia. Our data highlight the positive impact that transposable elements can have on the evolution of bacterial populations.

**Author summary:** Transposable elements naturally accumulate within genomes in all kingdoms of life. When present in the same orientation, a pair of homologous elements can act as substrates for DNA recombination reactions that can duplicate and delete intervening sequences – giving rise to genetically heterogenous populations. We showed here that *Burkholderia thailandensis* strain E264 uses this mechanism to amplify genes encoding a two-component regulatory system and an archaic chaperone usher pilus, priming the cells for rapid biofilm formation. The formation of a small subpopulation of biofilm-ready bacteria serves as a bet- hedging strategy, ensuring overall population survival should conditions change rapidly from those in which planktonic growth is optimal to those in which adherence and biofilm formation is required.

## Introduction

*Burkholderia thailandensis* is a saprophytic environmental bacterium native to tropical regions of Northern Australia and Southeastern Asia (1). *B. thailandensis* has evolved to survive in aquatic and soil environments, such as rice paddies, where it must adapt to fluctuations in environmental conditions and compete with other microbes.

As is true for organisms in all kingdoms of life, *B. thailandensis* has accumulated insertion sequences and transposons – parasitic genetic elements capable of moving between DNA sequences – throughout its genome (2). In *B. thailandensis* strain E264 (*Bt*E264) and close relatives, two nearly identical IS*2*-like elements, ISα and ISβ, bound a 208.6 kb region of DNA in chromosome I (hereafter referred to as ‘the 208.6 kb region’) (3). We recently discovered that these IS elements act as substrates for RecA-dependent homologous recombination that occurs during replication to produce tandem duplications of the 208.6 kb region (4). Reciprocally, homologous recombination between any of the duplicated 208.6 kb sequences in the tandem repeats can resolve the region back down to a single copy (4). Because these recombination reactions occur at a low but constant level, *Bt*E264 populations are consistently heterogeneous, composed of cells with (Dup+) and without (Dup-) duplicate copies of the 208.6 kb region (4).

Amplification of DNA sequences (increasing gene dosage) can alter cell phenotypes (5). A well-studied example is increased antibiotic resistance, which can occur by amplification of genes encoding antibiotic targets, efflux pumps, or antibiotic-modifying enzymes (6). Copy number of the 208.6 kb region influences phenotypes in *Bt*E264, impacting colony morphology, pigmentation, and most notably, the rate of biofilm formation (4). While Dup- and Dup+ *Bt*E264 can form visible biofilms on the sides on plastic culture tubes after multiple days of growth in M63 minimal medium, only Dup+ cells form visible biofilms at 24 hours post-inoculation – a rapid form of biofilm formation that we refer to as efficient biofilms (4).

Additionally, Dup+ cells have a competitive advantage in biofilms, whereas Dup- cells have a competitive advantage during planktonic growth (4). We previously posited that the generation of Dup+ and Dup- subpopulations is a bet-hedging strategy that allows *Bt*E264 to survive in unpredictable environments that quickly switch between those favoring biofilm or planktonic lifestyles (4).

The 208.6 kb region contains 158 predicted genes. We hypothesized that increasing the dosage of one or more of these genes leads to efficient biofilm formation. Our goals were to identify which gene(s) within the 208.6 kb region, when duplicated, confer this efficient biofilm phenotype, and to investigate the underlying mechanism.

## Results

### Duplicate copies of DNA containing genes predicted to encode an archaic-type chaperone-usher pilus, a small hypothetical protein, and a two-component regulatory system are sufficient to promote efficient biofilm formation

To identify the gene(s) responsible for efficient biofilm formation, we first divided the 208.6 kb region into nine subregions (Figure 1A), cloning each onto a plasmid using a modified plasmid rescue technique (Supplementary Figures 1 & 2). Briefly, we integrated a pair of I-*Sce*I restriction endonuclease site-containing suicide plasmids into the chromosome at the boundaries of each subregion, then isolated genomic DNA from each strain, digested the DNA with I-*Sce*I, ligated the isolated DNA, and transformed *E. coli*, selecting for kanamycin resistance (Km^R^). The composition and integrity of the resulting plasmids were confirmed by DNA sequence analysis. We then introduced each subregion-containing suicide plasmid into separate *Bt*E264 strains lacking ISβ (ΔISβ) (Supplementary Figure 3), which were therefore unable to duplicate the entire 208.6 kb region, resulting in strains in which only a single subregion was duplicated. We tested each strain for its ability to form biofilms after 24 hours growth in M63 medium. Only the strain with a duplicate copy of subregion 4 formed efficient biofilms (Figure 1A), indicating that the gene(s) sufficient for promoting efficient biofilm formation when present in multiple copies is/are within subregion 4.

**Figure 1.**
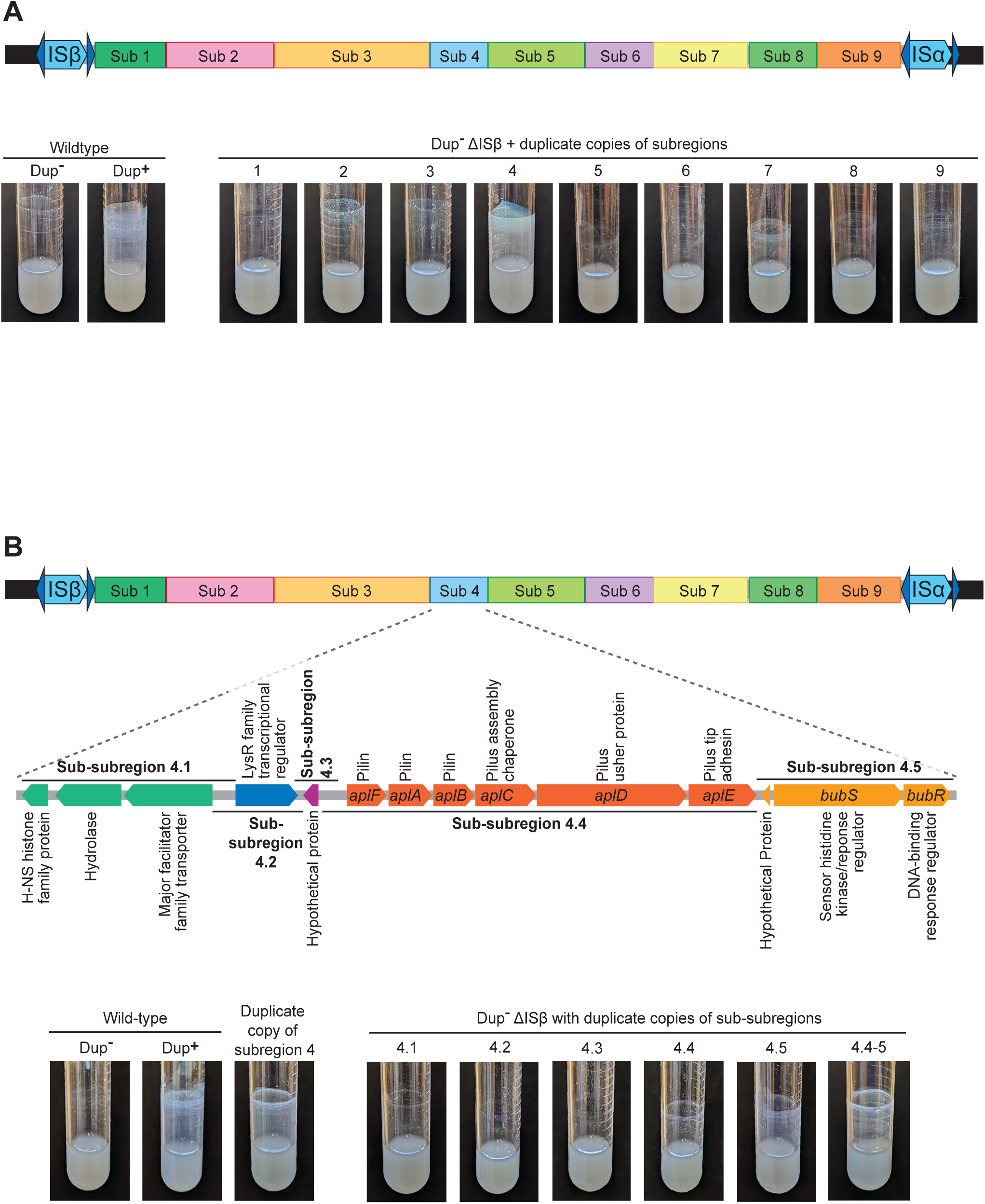
Legend: Duplicate copies of a subregion of DNA containing *aplFABCDE, iou, and bubSR,* is sufficient to promote efficient biofilm formation. (A) Top: Schematic of the duplicating region divided into nine subregions. Subregion boundaries were designed to prevent interruption of coding sequences or separation of coding sequences from their putative promoters. Bottom: Images of overnight cultures comparing the biofilm-forming ability of wild-type *B*tE264 to strains with duplicate copies of subregions 1 - 6. (B) Top: Schematic of the fourteen predicted genes located within subregion 4 and the boundaries of sub-subregions 4.1 – 4.5. Bottom: Images of overnight cultures comparing the biofilm-forming ability of wild-type *Bt*E264 and a strain with a duplicate copy of subregion 4 to strains with duplicate copies of sub- subregions 4.1 – 4.5 and 4.4-5.

Subregion 4 contains fourteen genes (Figure 1B). Thirteen predicted genes are annotated in the reference genome: one H-NS histone family protein, one hydrolase, one major facilitator family transporter, one LysR family transcriptional regulator, one protein of unknown function, six comprising an archaic-type chaperone-usher pilus system, one hypothetical protein, and two encoding a two-component regulatory system (TCS). Closer investigation of subregion 4 revealed an additional ORF between the pilus- and TCS-encoding operons that is oriented in the opposite direction and has the potential to encode a small protein of unknown function. We divided subregion 4 into five sub-subregions (Figure 1B), constructed suicide plasmids containing a single sub-subregion, and introduced these plasmids into the ΔISβ strain. Strains with a duplicate copy of sub-subregion 4.4 or 4.5 formed a minimal biofilm, but neither fully phenocopied the level of biofilm formed by strains with a duplication of the full 208.6 kb region or a duplication of subregion 4. However, integration of a plasmid containing both sub-subregions 4.4 and 4.5 together (sub- subregion 4.4-5), resulted in a strain that formed efficient biofilms (Figure 1B). We named the genes in sub- subregion 4.4-5 *aplFABCDE* for archaic pilus gene cluster, *iou* for intergenic orf of unknown function, and *bubSR* for *Burkholderia* biofilm sensor and regulator. Our data indicate that duplicate copies of the subregion of DNA containing these genes is sufficient for generating the efficient biofilm phenotype.

### *aplFABCDE* and *bubSR*, but not *iou*, are required for efficient biofilm formation

We described previously the development of ‘fragmented gene’ reporter strains, in which we replaced the IS*2*-like elements bounding the 208.6 kb region with a pair of incomplete, overlapping reporter gene fragments (4). The fragment replacing ISα is missing the 5ʹ end of the reporter gene and the fragment replacing ISβ is missing the 3ʹ end of the reporter gene. Homologous recombination within the overlapping central part of the gene fragments results in duplication of the region and, a functional copy of the reporter gene (conferring antibiotic resistance, fluorescence, or the production of β-glucuronidase) present at the junction between the two duplicated regions (4). These reporters allow us to select or screen for bacteria containing a duplication of the intervening sequences.

To determine if *aplFABCDE, iou*, or *bubSR* are required for efficient biofilm formation, we used the fragmented *nptII* (encoding kanamycin resistance (Km^R^)) reporter strain (Figure 2A). To mutate *aplFABCDE*, *iou*, and/or *bubSR*, we replaced them with *dhfRII* (encoding trimethoprim resistance (Tmp^R^)) in the fragmented *nptII* reporter strain, using bacteria that were Km^S^ and hence contained only a single copy of the region. We then selected Km^R^ derivatives (in which the region had duplicated) and compared Dup- (Km^S^) and Dup+ (Km^R^) bacteria for their ability to form efficient biofilms. The wild-type bacteria and the *iou* mutant formed similar biofilms after 24 hours, while the *aplFABCDE* and *bubSR* mutants failed to form biofilms (Figure 2B), indicating that *aplFABCDE* and *bubSR*, but not *iou*, are required for efficient biofilm formation.

**Figure 2.**
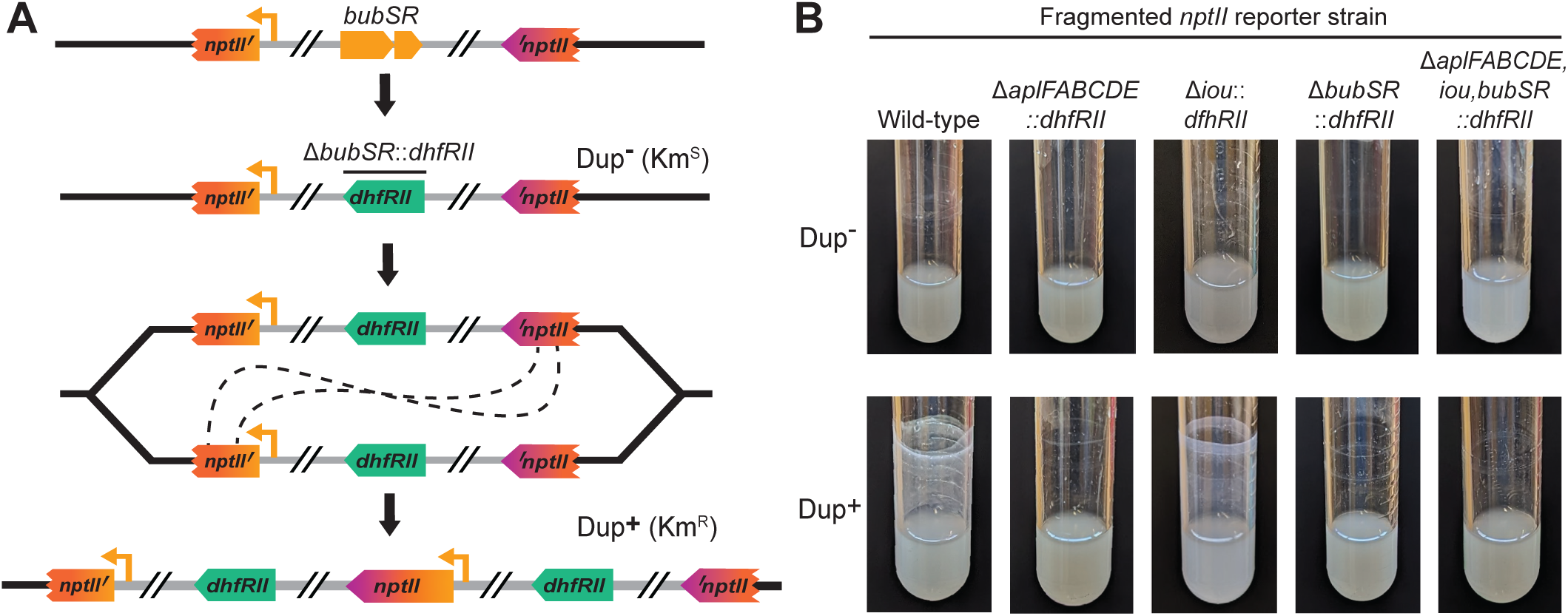
Legend: *aplFABCDE* and *bubSR* are required for efficient biofilm formation. (A) Schematic of the fragmented *nptII* gene system used to duplicate the 208.6 kb region in strains lacking sequences of interest, such as *bubSR*, to test requirements for efficient biofilm formation. Genes of interest were replaced with *dhfRII* in a Dup- (Km^S^) fragmented *nptII* reporter stain. Using kanamycin selection, cells that had spontaneously duplicated the region could be isolated. (B) Images of overnight cultures comparing the biofilm forming ability of fragmented *nptII* strains with deletions of *aplFABCDE*, *iou*, and/or *bubSR* with and without duplicate copies of the 208.6 kb region.

### *aplFABCDE* and *bubSR* are sufficient and required for duplication to provide a selective advantage during biofilm growth

We tested the hypothesis that *aplFABCDE* and *bubSR* are responsible for the selective advantage conferred by duplication of the 208.6 kb region during biofilm growth using our fragmented *gusA* (encoding β-glucuronidase) reporter system. With this system, Dup+ (*gusA*^+^) bacteria form blue colonies when plated on medium containing X-gluc. We replaced *aplFABCDE, iou*, and/or *bubSR* with *dhfRII* in Dup- fragmented *gusA* reporter bacteria then grew these strains to develop dynamic biofilms, replacing the liquid medium (and any cells it contained) every day with fresh, sterile medium. Throughout the experiment, mutants differed in their ability to form biofilms on the sides of the culture tube (Figure 3A). Wild-type and Δ*iou* strains formed visible biofilms within two days that increased in robustness as the experiment progressed. Meanwhile, Δ*aplFABCDE*::*dhfRII*, Δ*bubSR*::*dhfRII*, and Δ*aplFABCDE,iou,bubSR*::*dhfRII* strains were unable to form robust biofilms, and only formed minimal clumpy and nonuniform aggregations on the sides of the culture tube late into the experiment, which looked considerably different than the biofilms formed by wild-type and Δ*iou*::*dhfRII* bacteria. These data show that *aplFABCDE* and *bubSR* are not only required for efficient biofilm formation, but they are also required for longer-term ‘inefficient’ biofilm formation as well.

**Figure 3.**
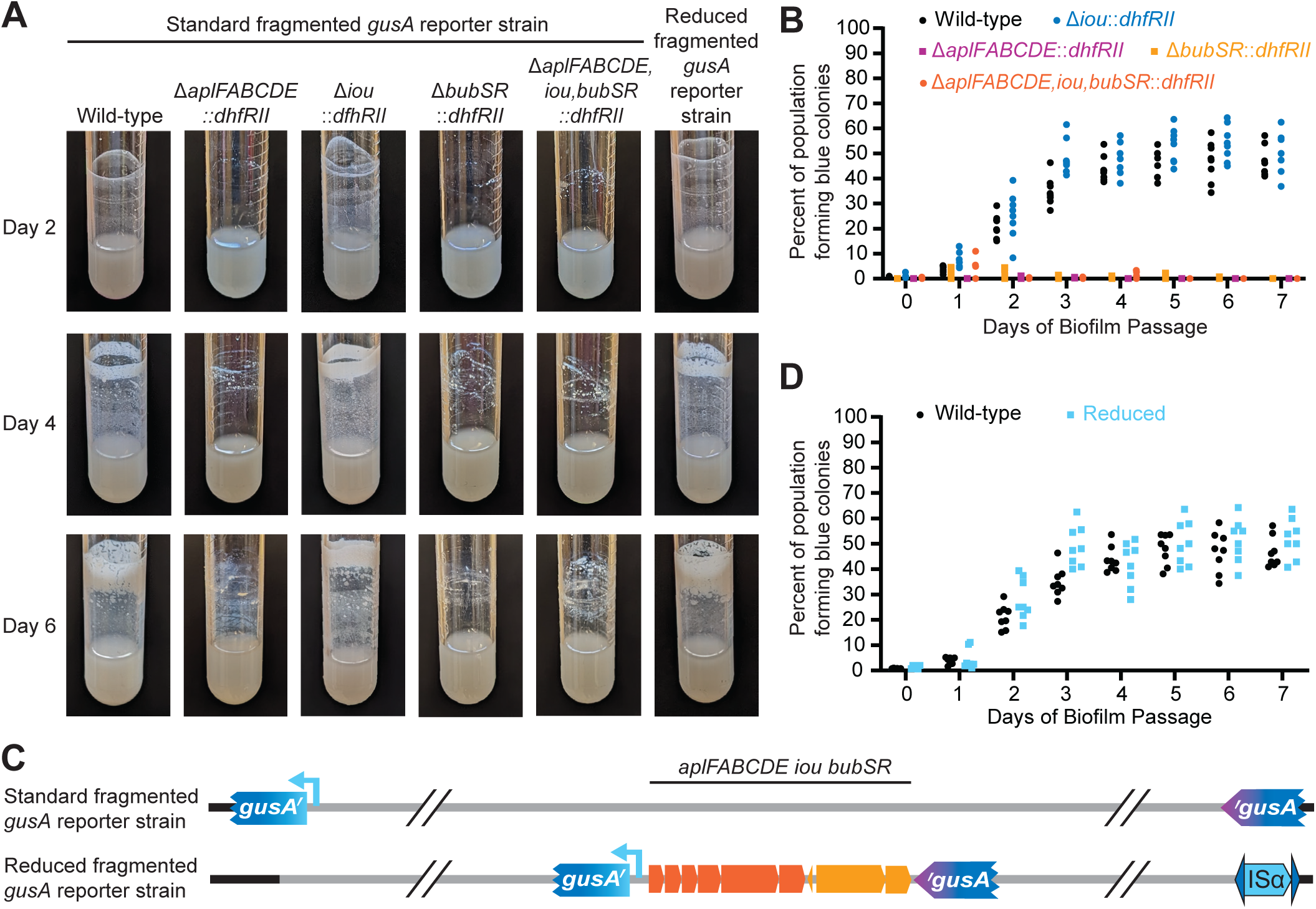
Legend: *aplFABCDE* and *bubSR* are sufficient and required for duplication to provide a selective advantage during biofilm growth. (A) Images of culture tube biofilms following two, four, and six days of serial biofilm passage of the fragmented *gusA* reporter strains that are otherwise wild-type or have *aplFABCDE*, *iou*, and/or *bubSR* deleted through replacement with *dhfRII.* Graph of the percentage of blue colonies grown from cells scraped from the edge of the culture tube daily during a seven-day serial biofilm passage experiment comparing (B) standard fragmented *gusA* reporter strains lacking *aplFABCDE, iou*, and/or *bubSR*, and (D) the standard and reduced fragmented *gusA* reporter strains. (C) Schematic of the fragmented *gusA* reporter strains wherein homologous recombination duplicates the entire 208.6 kb region (Standard) or just the 9.8 kb subregion containing *aplFABCDE,iou,bubSR* (Reduced).

When sampling from the biofilms (wild-type and Δ*iou::dhfRII* strains), we observed that within two days, nearly 25% of the population formed blue colonies on X-Gluc-containing medium, indicating that a quarter of the biofilm was composed of Dup+ cells (Figure 3B). At three days and longer, the percentage of blue colonies, and thus Dup+ cells, was nearly 50%. These data are consistent with those we reported for wild-type *Bt*E264 previously (4). These data indicate that *iou* is not required for the advantage that duplication of the region provides during biofilm growth (Figure 3B). For the Δ*aplFABCDE*::*dhfRII*, Δ*bubSR*::*dhfRII*, and Δ*aplFABCDE,iou,bubSR*::*dhfRII* strains, nearly all the colonies obtained from the scant ‘biofilms’ that formed were white on X-Gluc-containing medium, indicating that they were Dup-.a *plFABCDE* and *bubSR,* therefore, are required for the selective advantage conferred by duplication of the region during dynamic biofilm growth.

To determine if duplication of the *aplFABCDE,iou,bubSR* gene cluster is su>icient to confer the selective advantage, i.e., none of the remaining 149 genes within the 208.6 kb region are required, we generated a ‘reduced’ fragmented *gusA* reporter strain wherein we inserted the *gusA* gene fragments immediately flanking the *aplFABCDE,iou,bubSR*, genes (Figure 3C). When serially passaged to select for dynamic biofilm formation, the level and quality of biofilm formation mirrored that of the strain with *gusA* fragments at ISα and ISβ (Figure 3A). Similarly, when sampling from the biofilm, the proportion of blue colonies increased and plateaued at nearly 50% after three days (Figure 3D). These data indicate that duplication of the *aplFABCDE,iou,bubSR* gene cluster is su>icient for providing the selective advantage during biofilm growth.

### BubSR TCS activity is required for e<icient biofilm formation

Analysis of the predicted amino acid sequence of BubS with SMART domain prediction and AlphaFold2 reveals that BubS is likely an unorthodox histidine kinase (i.e., it contains a histidine kinase domain, a receiver domain, and a histidine phosphotransfer domain) (Figure 4A). It is unusual in that it contains no obvious sensory input domain. Histidine Kinase A (dimerization/phosphoacceptor) (HisKA) (BubSA84 – G149) and Histidine Kinase-like ATPase (HATPase) (BubST196-V312) domains were identified by SMART, and the predicted phosphorylation site is H94. A CheY-similar Receiver (REC) domain (BubSY436-L551) was also predicted by SMART, with conserved aspartic acid residues D442, D443, and D486: D486 being the site of phosphorylation. No Histidine Phosphotransfer (HPt) domain was predicted by SMART, but AlphaFold2 analysis of the C-terminal 150 amino acids of BubS (BubSE556-S680) revealed a high-confidence structure that resembles the Hpt of the *Escherichia coli* unorthodox histidine kinase BarA (Figure 4B and C).

**Figure 4.**
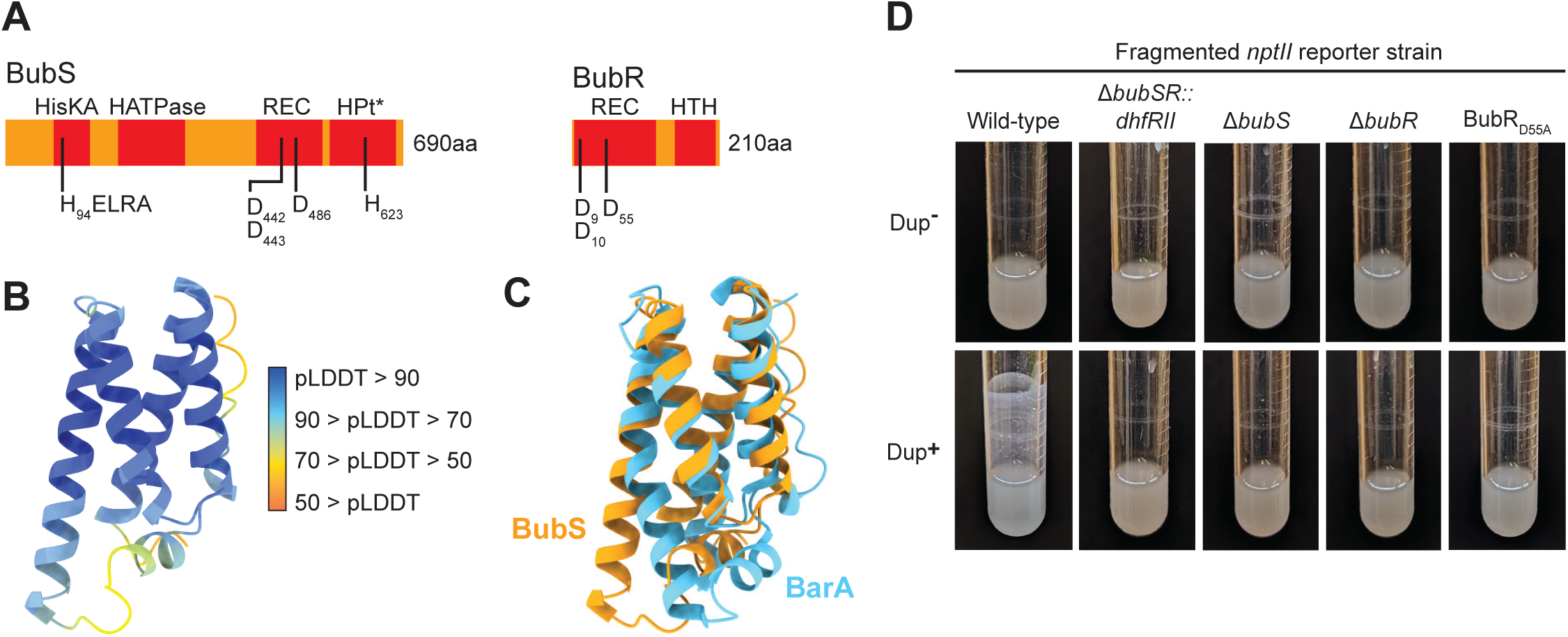
Legend: BubSR TCS activity is required for e<icient biofilm formation. (A) Schematic of the BubS and BubR proteins including predicted domains identified by SMART and AlphaFold2 (denoted with *). Amino acids involved in phosphoryl relay are denoted. (B) AlphaFold2 structure of BubSE556-S680. The structure is colored according to per-residue predicted local distance di>erence test (pLDDT) scores that quantify structure confidence. (C) AlphaFold2 structure of BubSE556-S680 overlaid with the *E. coli* BarA HPt domain (3IQT). (D) Images of overnight cultures comparing the biofilm forming ability of wild-type and *bubSR*::*dhfRII*, Δ*bubS*, Δ*bubR*, and BubRD55A mutants in a fragmented *nptII* background strain with and without duplicate copies of the 208.6 kb region.

SMART predicted REC and LuxR-type Helix-turn-helix DNA binding (HTH) domains in BubR, indicating that BubR belongs in the NarL/FixJ family. The BubR REC domain likely coordinates the phosphoryl group at residues D9, D10, and D55, with D55 as the site of phosphorylation.

To determine if BubSR TCS activity is required for Dup+ cells to produce e>icient biofilms, we used allelic exchange in a fragmented *nptII* reporter strain to delete *bubS* or *bubR* or to change codon 55 in *bubR* such that it encodes alanine instead of aspartic acid (BubRD55A), which should prevent phosphorylation of BubR. All three mutations, regardless of copy number of the region, abrogated the ability of the bacteria to form e>icient biofilms (Figure 4D), indicating that BubSR activity is required for e>icient biofilm formation.

### Two copies of *bubSR* plus the intergenic region 5ʹ to *aplFABCDE* are sufficient to promote efficient biofilm formation

Because wild-type *Bt*E264 and *bubSR* mutants are indistinguishable when grown planktonically in M63 medium, but only wild-type (not *bubSR* mutant) bacteria can form biofilms (either in 24 hours for Dup+ bacteria or four days for Dup-bacteria), we reasoned that planktonic growth in M63 medium is likely a condition in which the BubSR TCS is inactive, and biofilm growth is likely a condition in which BubSR is active. We hypothesized that *bubSR* is positively autoregulated and that active BubR (i.e., phosphorylated BubR (BubR∼P)) also activates transcription of the *aplFABCDE* genes, which are also required for biofilm formation. Moreover, we hypothesized that two copies of *bubSR* causes the level of BubR∼P in at least some bacteria growing planktonically in M63 to be above the threshold needed for positive autoregulation. To test this hypothesis, we cloned *bubSR* plus the intergenic region between *aplE* and *bubS* (the putative promoter region for *bubSR*) into plasmid pUC18-miniTn7-Km and delivered the resulting plasmid into a Δ*bubSR::dhfRII* ΔISα strain by conjugation. Because *B. thailandensis* E264 contains two *glmS* genes, we could obtain strains containing *bubSR* (and the 3ʹ adjacent *nptII* gene, which is not shown in the figure) inserted at either one or both *att*Tn*7* sites (Figure 5A). Unexpectedly, the strain containing two copies of *bubSR* (one at each *att*Tn*7* site) did not form efficient biofilms (Figure 5A). We next cloned the region containing the entire *aplFABCDE,iou,bubSR* locus, including the intergenic region 5ʹ to *aplF,* into plasmid pUC18-miniTn7-Km and selected strains in which the *aplFABCDE,iou,bubSR* (and *nptII*, not shown)- containing cassette was present at one or both *att*Tn7 sites in a Δ*aplFABCDE,iou,bubSR*::*dhfRII* ΔISα strain (Figure 5B). Only the strain containing two copies of the *aplFABCDE,iou,bubSR* locus formed an efficient biofilm (Figure 5B). It seemed unlikely to us that two copies of *aplFABCDE* are required for efficient biofilm formation. Instead, we hypothesized that the promoter required for activating transcription of *bubSR* is located 5ʹ to *aplF*. To test this hypothesis, we constructed a plasmid containing the entire *aplFABCDE,iou,bubSR* locus but with a large deletion extending from *aplF* to *aplE,* and delivered the plasmid to the Δ*bubSR::dhfRII* ΔISα strain by conjugation (Figure 5C). Surprisingly, even the strain with the *aplF*ʹ*-*ʹ*aplE,iou,bubSR-*containing cassette at only one *att*Tn*7* site formed an efficient biofilm (Figure 5C).

**Figure 5.**
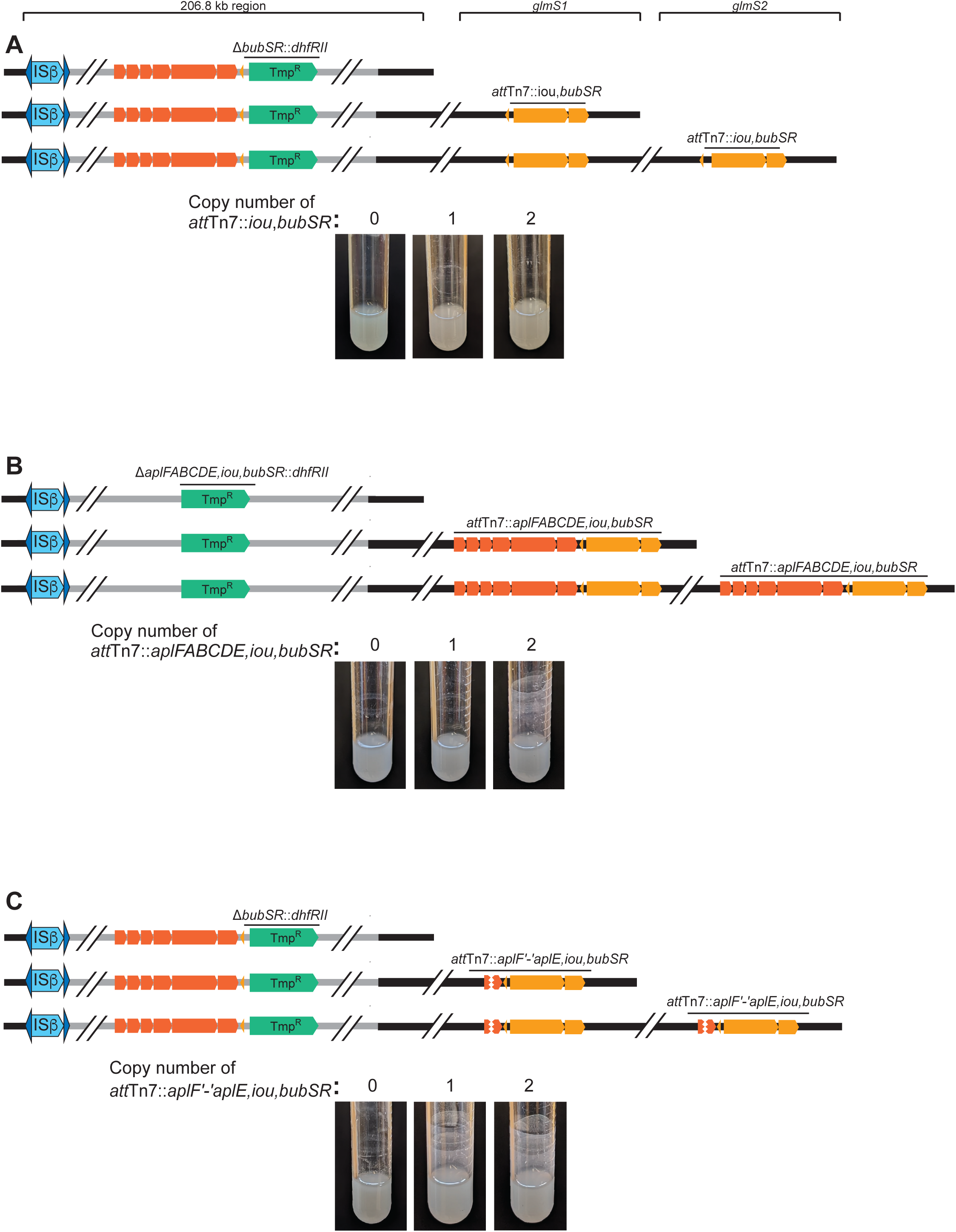
Legend: Two copies of *bubSR* plus the intergenic region 5ʹ to *aplFABCDE* are sufficient to promote efficient biofilm formation. (A) Top: Schematic of strains with *bubSR* replaced with *dhfRII* and with *bubSR* inserted at either one, both, or neither *att*Tn*7* site. Bottom: Images of overnight cultures. (B) Top: Schematic of strains with *aplFABCDE,iou,bubSR* replaced with *dhfRII* and with *aplFABCDE,iou,bubSR* inserted at either one, both, or neither *att*Tn*7* site. Bottom: Images of overnight cultures. (C) Top: Schematic of strains with *bubSR* replaced with *dhfRII* and with *aplF*ʹ*-*ʹ*aplE,iou,bubSR* inserted at either one, both, or neither *att*Tn*7* site. Bottom: Images of overnight cultures.

These results indicate that two copies of the *aplFABCDE* genes are not required for efficient biofilm formation, and they suggest that a promoter required for activation of both *aplFABCDE* and *bubSR* expression is located in the intergenic region 5ʹ to *aplF*. The data also suggest that sequences within the *aplFABCDE* genes somehow negatively influence *bubSR* expression.

### The intergenic region 5ʹ to *aplF*, but not the region between *aplE* and *bubS*, contains a promoter that is activated during biofilm growth and when the 208.6 kb region is duplicated, in a *bubSR*-dependent manner

To determine directly if the regions 5ʹ to *aplF* and *bubS* contain promoters that are activated when the 208.6 kb region is duplicated and/or during biofilm growth, we cloned each putative promoter region (P_*apl*_ and P_*bub*_, Figure 6A) 5ʹ to the *gfp* gene that was codon-optimized for *B. thailandensis* in plasmid pKBM16 (creating pKBM21 and pKBM20, respectively), and delivered the P*apl-gfp* and P*bub-gfp* cassettes to *att*Tn*7* (selecting those with only a single copy at the *glmS1*site) in wild-type and *bubSR* mutant strains containing either a deletion of ISα (Dup- locked bacteria) or the fragmented *nptII* genes (to select for Dup+ bacteria). For the resulting GFP reporter strains, we grew cultures wherein spent medium was replaced with fresh M63 every 24 hours. Planktonic and biofilm bacteria were collected after 24- or 96-hours growth in M63 medium, washed, fixed, and transferred to a 96-well plate to measure fluorescence intensity.

**Figure 6.**
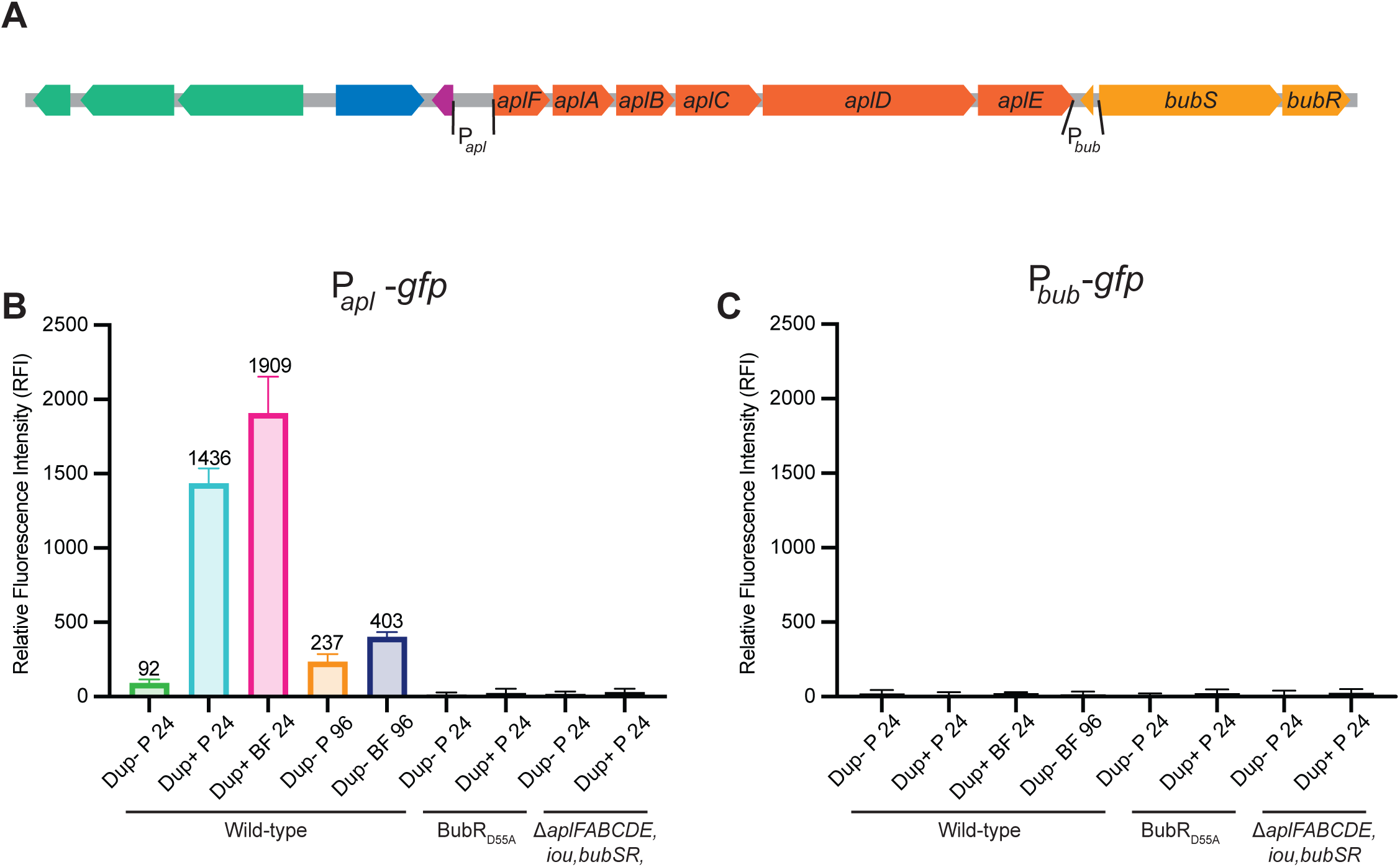
Legend: The intergenic region 5ʹ to *aplF*, but not the region between *aplE* and *bubS*, contains a promoter that is activated during biofilm growth and when the 208.6 kb region is duplicated, in a *bubSR*-dependent manner. (A) Schematic of subregion 4 with putative *apl* and *bub* promoter regions used for construction of promoter-*gfp* fusions indicated. (B) Relative fluorescence intensity of the indicated strains containing the P_*apl*_-*gfp* fusion after 24- or 66-hours growth. P = planktonic, BF = biofilm. (C) Relative fluorescence intensity of the indicated strains containing the P_*bub*_-*gfp* fusion after 24- or 66- hours growth. P = planktonic, BF = biofilm.

Planktonically grown cells containing the P*apl-gfp* fusion produced a low level of fluorescence (62 RFI units) when the 208.6 kb region was present in a single copy after 24 hours growth in M63. By contrast, when the 208.6 kb region was duplicated, the level of fluorescence in bacteria growing planktonically was 1,436 RFI units, a 15.6-fold increase compared to Dup- bacteria (Figure 6B). When Dup+ bacteria containing the P*apl-gfp* fusion were recovered from the biofilm after 24 hours growth, the level of fluorescence was 1,606 RFI units (Figure 6B). Fluorescence of *bubS* or *bubR* mutants containing the P*apl- gfp* fusion was at background levels under all conditions (Figure 6B), confirming that activation of P_*apl*_ requires active BubSR.

In Dup- bacteria containing the P*apl-gfp* fusion grown for 66 hours, the level of fluorescence was 237 RFI units in the planktonic bacteria and 403 RFI units in the bacteria recovered from the biofilm (Figure 6C).

Strains containing the P*bub-gfp* fusion produced no fluorescence under any condition, confirming that the region 5ʹ to *bubS* does not contain a promoter that is active under any of the growth conditions tested (Figure 6C).

These data show that there is a promoter 5ʹ to *aplF* and that its expression increases dramatically, in a BubSR-dependent manner, when the 208.6 kb region is duplicated, suggesting that *bubSR* is positively autoregulated via P_*apl*_. P_*apl*_ is also activated during biofilm growth in Dup- bacteria, although not as strongly as in Dup+ bacteria.

### Duplication of *bubSR* is sufficient to activate P_*apl*_ in a majority of bacteria growing in M63 medium

The plate reader (data shown in Figure 6) measures total fluorescence in each population of bacteria. To measure fluorescence, and hence P_*apl*_ and P_*bub*_ expression, in single cells, we analyzed aliquots of the same cultures using flow cytometry. We stained the samples with SYTO 61 to identify cells and to exclude any debris and electronic background. Next, we used forward and side scatter to gate for single bacterial cells and measured the GFP fluorescence of particles contained within these parameters. The level of fluorescence for all bacteria containing the P_*bub*_-*gfp* fusion was very low (Figure 7B), as expected.

**Figure 7.**
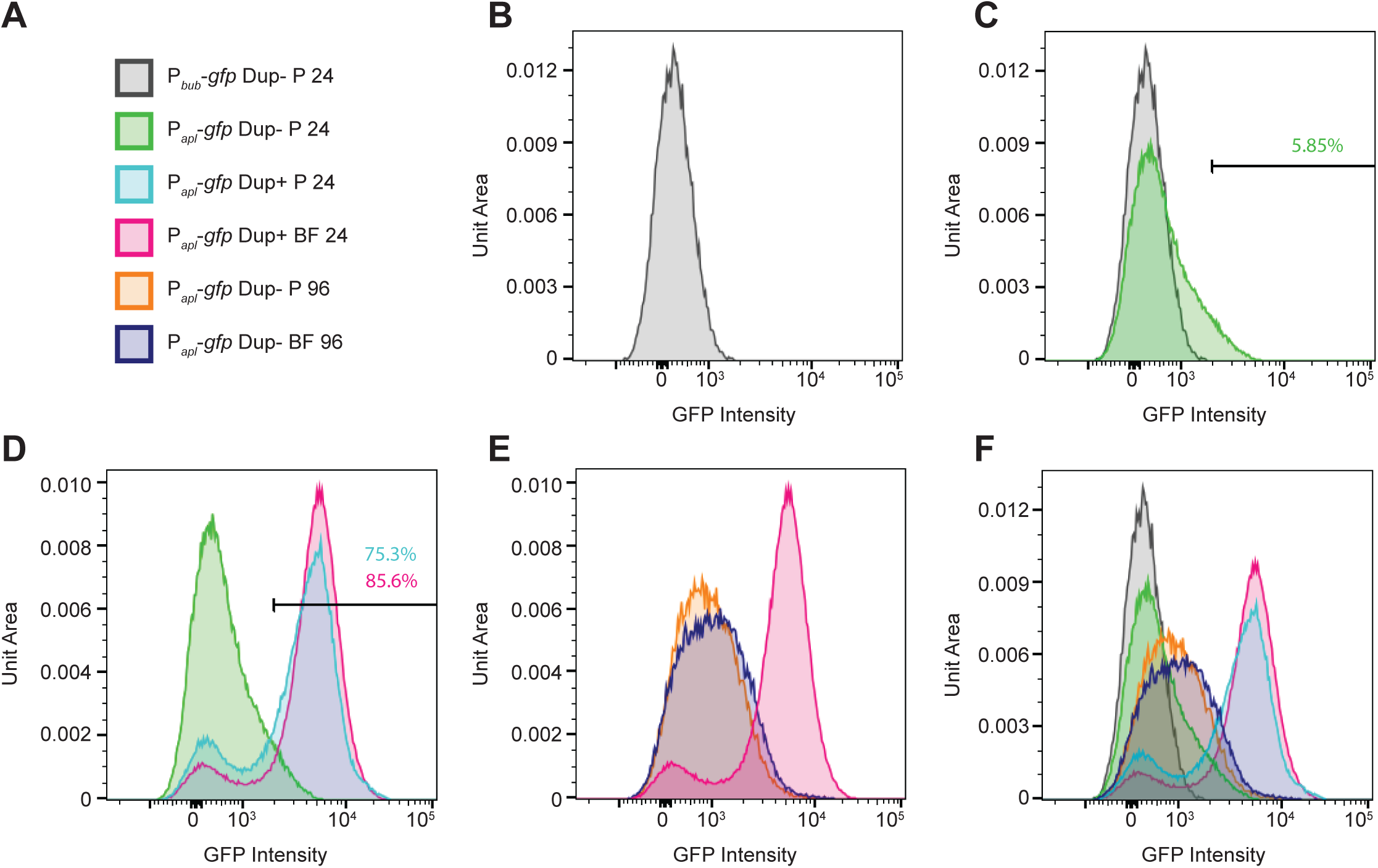
Legend: Duplication of *bubSR* is sufficient for positive autoregulation in a majority of bacteria growing in M63 medium. (A) Legend for the graphs shown in B–F. P = planktonic, BF = biofilm. (B-F) Graphs showing fluorescence of populations of bacteria as indicated by the legend. The level of fluorescence is plotted along the X-axis on a log scale, and the number of bacteria, normalized to unit area under each curve so that different samples can be compared, is plotted along the Y-axis.

For most Dup- planktonic bacteria containing the P*apl-gfp* fusion, the level of fluorescence was also very low, similar to that of bacteria containing the P_*bub*_-*gfp* fusion (Figure 7C, green curve). However, the level of fluorescence was higher in ∼6% of these bacteria and was quite high in a very small number of bacteria (Figure 7C, shoulder of green curve). These data are consistent with planktonic growth corresponding to a BubSR-inactive condition, with a small proportion of bacteria stochastically producing enough BubR∼P to activate P_*apl*_, and hence cause positive autoregulation of *bubSR*.

For Dup+ planktonic bacteria containing the P*apl-gfp* fusion, two distinct populations were apparent (Figure 7D, cyan curve): 24.7% of the population was essentially non-fluorescent and 75.3% of the population was highly fluorescent, indicating that two copies of *bubSR* is sufficient for activation of P_*apl*_ – and hence also *bubSR* – even under BubSR-inactivating conditions in most bacteria. Dup+ bacteria containing the P*apl-gfp* fusion collected from the biofilm (Figure 7D, pink graph) had an even higher proportion of highly fluorescent bacteria (85.6%). Dup- bacteria containing the P*apl-gfp* fusion collected after 66-hours growth displayed a range of fluorescence from low to moderate, with a slight shift towards increased fluorescence in bacteria collected from the biofilm compared to bacteria recovered from the liquid (Figure 7E). Although very few of these bacteria were as fluorescent as Dup+ bacteria.

Together, these data suggest that planktonic growth in M63 medium is a condition in which the BubSR TCS is inactive, but that a small amount of BubR∼P forms stochastically in a small proportion of bacteria under these conditions, and that this level of BubR∼P is sufficient to activate P_*apl*_, and hence cause positive autoregulation of *bubSR*, which increases the amount of BubR∼P in those cells even further. The data suggest further that simply doubling the copy number of *bubSR* causes the level of BubR∼P that forms stochastically to be above the threshold required for activation of P_*apl*_ in a majority (∼75%) of bacteria.

## Discussion

To persist, bacterial populations must adapt to ever-changing environmental conditions, and they frequently do so using two-component regulatory systems (TCS). Composed generally of a histidine kinase that senses environmental cues and, in response, phosphorylates a response regulator that effects a change in behavior (usually by activating and/or repressing gene expression), TCSs allow all members of a population to adapt appropriately, maximizing their fitness to a specific environmental condition (7).

However, when facing unrecognizable stimuli, or when conditions fluctuate too rapidly for signal transduction systems to mount an appropriate response, bacterial populations may be unable to adapt. In these cases, bet-hedging strategies, wherein a subpopulation of bacteria stochastically exhibits a phenotype that is maladapted for the current condition but essential in another, may allow population survival (8). Bet-hedging can be accomplished by heritable and reversible genetic mutations that lead to stochastic switching between distinct phenotypes, a phenomenon known as phase variation (6). It can also be accomplished by unequal activation of signal transduction systems in genotypically homogenous bacterial populations, resulting in subpopulations that display phenotypes opposite to those dictated by the environmental condition (10,11).

We showed in this study that a combination of these two strategies – deterministic two- component signal transduction and stochastic, duplication-mediated activation of that TCS – underlies efficient biofilm formation in *B. thailandensis* E264. Our data showing that *bubSR* mutants are indistinguishable from wild-type bacteria during planktonic growth and are unable to form biofilms strongly suggest that the BubSR TCS is inactive during planktonic growth in M63 liquid medium and active during biofilm growth. This conclusion is supported by the promoter-*gfp* fusion data that indicate that in Dup- bacteria growing planktonically, P_*apl*_ expression, which is dependent on BubSR activity (and specifically BubR∼P, as the BubRD55A mutant fails to activate P_*apl*_), is very low in ∼94% of the bacteria. When Dup- bacteria are recovered from the inefficient biofilm that ultimately forms, P_*apl*_ expression is ∼4-fold higher than in planktonic bacteria, indicating that BubSR is active in the biofilm. Thus, BubSR appears to mediate the deterministic response, controlling expression of at least two sets of genes (*aplFABCDE* and *bubSR*) that are required for biofilm growth, such that they are expressed under biofilm conditions and not during planktonic growth. Stochastic, RecA-dependent duplication of the 208.6 kb region provides the bet-hedging strategy; in 75% of Dup+ bacteria growing planktonically P_*apl*_ expression was very high, indicating that simply doubling the amount of BubSR causes the level of BubR∼P, under BubSR-inactivating conditions, to be above the threshold required to activate P_*apl*_, resulting in greatly increased expression of the *aplFABCDE* and *bubSR* genes. In wild-type bacteria growing planktonically (such as, perhaps, in flooded soil), the proportion of Dup+ bacteria is likely to be about one in ∼10,000 (4), so even if 75% of those bacteria have an active BubSR TCS, the proportion of maladapted bacteria in the population will be very small. However, if conditions change rapidly such that biofilm growth is required, the small proportion of BubSR-active bacteria will survive, maintaining the lineage at that site. Thus, the bet-hedging strategy provided by RecA-dependent duplication of the 208.6 kb region may allow population survival when conditions shift from those favoring planktonic growth to those requiring biofilm growth too rapidly for a BubSR signal transduction-mediated response.

For TCSs to function, there must be sufficient sensor kinase and response regulator proteins in the cell under ‘OFF’ conditions for the sensor kinase to be able to respond to ‘ON’ conditions by autophosphorylating and transferring the phosphoryl group to, and thus activating, the response regulator, which, in many cases, causes positive autoregulation and amplification of the system. Simple enzyme kinetics would predict that even under ‘OFF’ conditions, a small proportion of response regulator protein would be phosphorylated stochastically. Our data suggest that for BubSR, the proportion of planktonic bacteria after overnight growth in M63 with enough BubR∼P to activate P_*apl*_ is about 6%, which is higher than we would have predicted. Moreover, simply doubling *bubSR* copy number increased that proportion to ∼75%. These data suggest that BubS activity may be just below an activation threshold when the bacteria are growing in M63 medium such that merely doubling the amount of BubS pushes the system over the threshold in most cells. The lack of obvious sensing domains makes predicting the signals to which BubS responds difficult. However, *bubSR* homologs have been identified in *Burkholderia pseudomallei* (12,13), and there is evidence that *bubSR* (called *bfmSR* in *B. pseudomallei*) expression and BubSR (BfmSR) activity may increase under iron-limiting conditions (14). It is possible that after overnight growth in M63 medium, and maybe more so for bacteria accumulating on the walls of the test tube, iron and/or other nutrients are depleted, shifting the conditions towards those in which BubS is active. We will investigate this possibility in the future.

Comparisons between *B. thailandensis* and *B. pseudomallei* yielded additional insight. Although the amino acid sequences of BubS in *Bt*E264 and BfmS in *B. pseudomallei* K66243 are 60% identical and 62% similar, SMART predicts a transmembrane domain at the N-terminus of BfmS but not for BubS. As most sensor kinases are cytoplasmic membrane proteins, it is likely that BubS contains a trans-membrane domain that was not identified by SMART, but biochemical experiments will be required to determine if BubS (or BfmS) is membrane-localized. It is also noteworthy that although the intergenic sequences between *aplFABCDE* (*B. thailandensis*)/*csuABCDE* (*B. pseudomallei*) *and bubS* (*B. thailandensis*)*/bfmS* (*B. pseudomallei*) are nearly identical, a single additional nucleotide in the *B. pseudomallei* sequence abrogates the prediction of an ORF in this region. This information, together with the fact that *iou* is not required for either efficient or inefficient biofilm formation or for the selective advantage conferred by duplicating the 208.6 kb region during biofilm growth in *Bt*E264, suggests that *iou* may not encode a functional protein.

Genetic linkage between genes encoding regulatory proteins and the genes they control is common, and it is also common for TCSs to (also) regulate unlinked genes. While the full BubSR regulon is currently unknown, genes in addition to *aplFABCDE*, including those predicted to encode an exopolysaccharide, are required for biofilm formation (15). In a previous report from our group, we compared the transcriptomes of wild-type *Bt*E264 with those of strains in which expression of the *bcpAOIB* genes (encoding a contact-dependent inhibition system) was driven by the constitutive S12 promoter or that produced a catalytically inactive BcpA protein (15). We now know that these strains were predominantly Dup+ and Dup-, respectively. Consistent with our current data, transcript abundance for the *aplFABCDE* and *bubSR* genes was dramatically increased in the strain with PS12 driving *bcpAIOB* compared to the BcpA mutant, as were many others, including those predicted to encode an exopolysaccharide. However, because the 208.6 kb region contains several genes that are predicted to encode transcription regulators, additional experiments will be required to determine if any of the differentially-expressed genes identified in the previous study are controlled by BubSR.

In the current study, we focused on efficient biofilm formation, but other duplication-dependent phenotypes, such as Congo red binding and the production of a gold-brown pigment, exist (3,4). We do not know, at this point, what genes are responsible for those phenotypes or whether they are regulated by BubSR. Indeed, these other phenotypes could require duplication of a different subregion within the 208.6 kb region. If these other duplication-dependent phenotypes are beneficial, it could explain the selective advantage for strains with ISα and ISβ in their current locations. We showed in this work that duplication of only the *aplFABCDE,iou,bubSR* genes is sufficient for efficient biofilms to form and for the selective advantage conferred by duplication during biofilm growth. Since duplication of a smaller region would provide less homology for recombination-mediated loss of the duplication, it would seem to be more advantageous for ISα and ISβ to be closer together. Are their current locations just chance, or is duplication of other genes within the 208.6 kb region advantageous under conditions that we have not yet explored? Continued investigation will be required to understand the depth and breadth of the stochastic, IS-mediated bet-hedging strategy in *Bt*E264.

Interplay between TCS signaling and phase variation is not uncommon. In many cases, phase variation removes one or more genes from the deterministic control of the TCS without altering the entire regulon. For example, phase variation-mediated mutation of genes encoding *Helicobacter pylori* adhesins and *Neisseria meningitidis* hemoglobin receptors, which are regulated by TCSs that control many genes required for infection, prevents production of these immunostimulatory proteins, thereby allowing the bacteria to evade the host immune system (16–20). In other cases, phase variation toggles on or off production of the TCSs themselves, leading to two distinct subpopulations; phase variation-ON subpopulations that produce the TCS and are able to sense-and-respond to the recognized stimulus, and phase variation-OFF subpopulations that are effectively blind to the activating stimulus (21–26). DNA duplication-mediated activation of BubSR represents another form of interplay between TCS signaling and phase variation, further highlighting the positive impact that transposable elements can have on the evolution of bacterial populations.

## Materials and Methods

### Bacterial culture conditions

*Bt*E264 is an environmental isolate (1). Plasmids were maintained in *E. coli* DH5α. For insertion at the *att*Tn*7* sites or Flp-mediated FRT recombination, plasmids were introduced into *Bt*E264 by conjugation with *E. coli* donor strain, RHO3 (27). *Bt*E264 and *E. coli* strains were grown overnight with aeration at 37°C (unless indicated) in low-salt Luria-Bertani (LSLB, 0.5% NaCl). Antibiotics and supplements were added to cultures at the following concentrations: 50 μg/mL X-Gluc (5-bromo-4- chloro-3-indoxyl-beta-D-glucuronide), 200 μg/mL 2,6-diaminopimelic acid (DAP), 0.2% (wt/vol) rhamnose, 500 μg/mL (for *Bt*E264) or 50 μg/mL (for *E. coli*) kanamycin (Km), 100 μg/mL (for *Bt*E264) or 50 μg/mL (for *E. coli*) trimethoprim (Tmp), 50 μg/mL (for *Bt*E264) or 10 μg/mL (for *E. coli*) tetracycline (Tc), 100 μg/mL ampicillin (Ap), 200 μg/mL zeocin (Zeo), or 30 μg/mL (for *Bt*E264) chloramphenicol (Cm) as appropriate. Because trimethoprim exposure alters gene expression in *Burkholderia thailandensis*, trimethoprim was only used to select for stable mutations that do not require continuous selection, and experiments were conducted without trimethoprim (28). When indicated, *Bt*E264 was cultured in M63 minimal medium (110 mM KH2PO4, 200 mM K2HPO4, 75 mM (NH4)2SO4, 16 nM FeSO4) supplemented with 1 mM MgSO4, 0.2% glucose, 0.4% glycerol, and 0.01% casamino acids.

### Mutant construction techniques

#### Natural transformation

Linearized plasmids containing an antibiotic resistance-encoding gene flanked by ∼500 bp sequences with homology to genomic regions of interest were introduced to *Bt*E264 following previously described protocols (29). Transformants were isolated on LSLB-supplemented with the appropriate antibiotic and verified through PCR analysis.

#### Allelic exchange

Markerless mutations were constructed through allelic exchange with *sacB* counterselection. Plasmids with a pEXKm5 backbone were constructed to contain mutant DNA sequences with homology to the chromosome and subsequently introduced into *E. coli* RHO3. RHO3 were then mated with a *Bt*E264 strain of interest and transformants were isolated on LSLB-supplemented with the appropriate antibiotic. Allelic exchange was conducted using previously established protocols (27).

#### Plasmid integration

Plasmids containing sequences of interest with homology to the genome were introduced into RHO3 and were subsequently mated into *Bt*E264, in which, the plasmid backbone could not replicate. Single homologous recombination between the chromosome and the introduced plasmid would integrate plasmid DNA onto the chromosome. Transformants were isolated on LSLB-supplemented with the appropriate antibiotic.

#### Flp-FRT recombination

Flp-mediated FRT recombination was used to excise antibiotic cassettes flanked by FRT sequences. Recombination was conducted according to previously described protocols (30).

#### attTn7 site Insertion

Introduction of DNA sequences to one or both *att*Tn7 sites was conducted according to previously described protocols (31) through mating between *Bt*E264, *E. coli* RHO3 carrying the Tn7 transposase- containing plasmid pTNS3, and *E. coli* RHO3 carrying a plasmid with DNA sequences for insertion. Mutants were confirmed with PCR analysis.

### Efficient Biofilm Growth

Overnight cultures in LSLB were diluted to an OD600 of 0.2 in 2ml of M63 within 14 mL polystyrene test tubes. Cultures were grown on a rotator at 37°C for 24 hours before imaging.

### Dynamic Biofilm Selection

Dynamic biofilm selection through serial passaging was conducted as described previously (4). Populations were initially composed of 2% Dup+ bacteria.

### Plate Reader Assay

Reporter strains were grown overnight in 2 mL LSLB at 37°C. Each culture was then diluted to 0.1 OD600 in 2 mL M63 in polystyrene tubes and grown at 37°C. All planktonic cells and Dup+ cells contained within a biofilm were collected after 24 hours of growth. Strains locked as Dup- were passaged daily with 2 mL M63 for 96 hours, at which point planktonic cells and cells contained within a biofilm were collected. At the time of collection, cells contained within a biofilm were washed o> the sides of the tubes with 1 mL PBS. All cells were then washed in PBS twice and fixed with 2% paraformaldehyde.

A clear 96-well plate was used to measure the OD600, and a black 96-well plate was used to measure the absolute fluorescence intensity (AFI) at ex. 485 nm, em. 535 nm. Planktonic cells were added to each plate at a 1:10 dilution in PBS to a final volume of 200 μL, and 200 μL of cells recovered from the biofilm were added to each plate. The OD600 and AFI of PBS were subtracted from the recorded values of each sample. The relative fluorescence intensities (RFI) were calculated by dividing the normalized AFI by the normalized OD600 for each sample.

### Flow Cytometry

Fixed cells used in the plate reader assay were subsequently analyzed using flow cytometry. Each sample was stained with 0.05 mM SYTO^TM^ 61 red fluorescent nucleic acid stain to distinguish cells from electronic noise and debris within the sample. Events positive for SYTO^TM^ 61 were then used to determine the gating parameters for forward and side scatter to identify single cells. Analysis was performed with FlowJo software.

## Plasmids

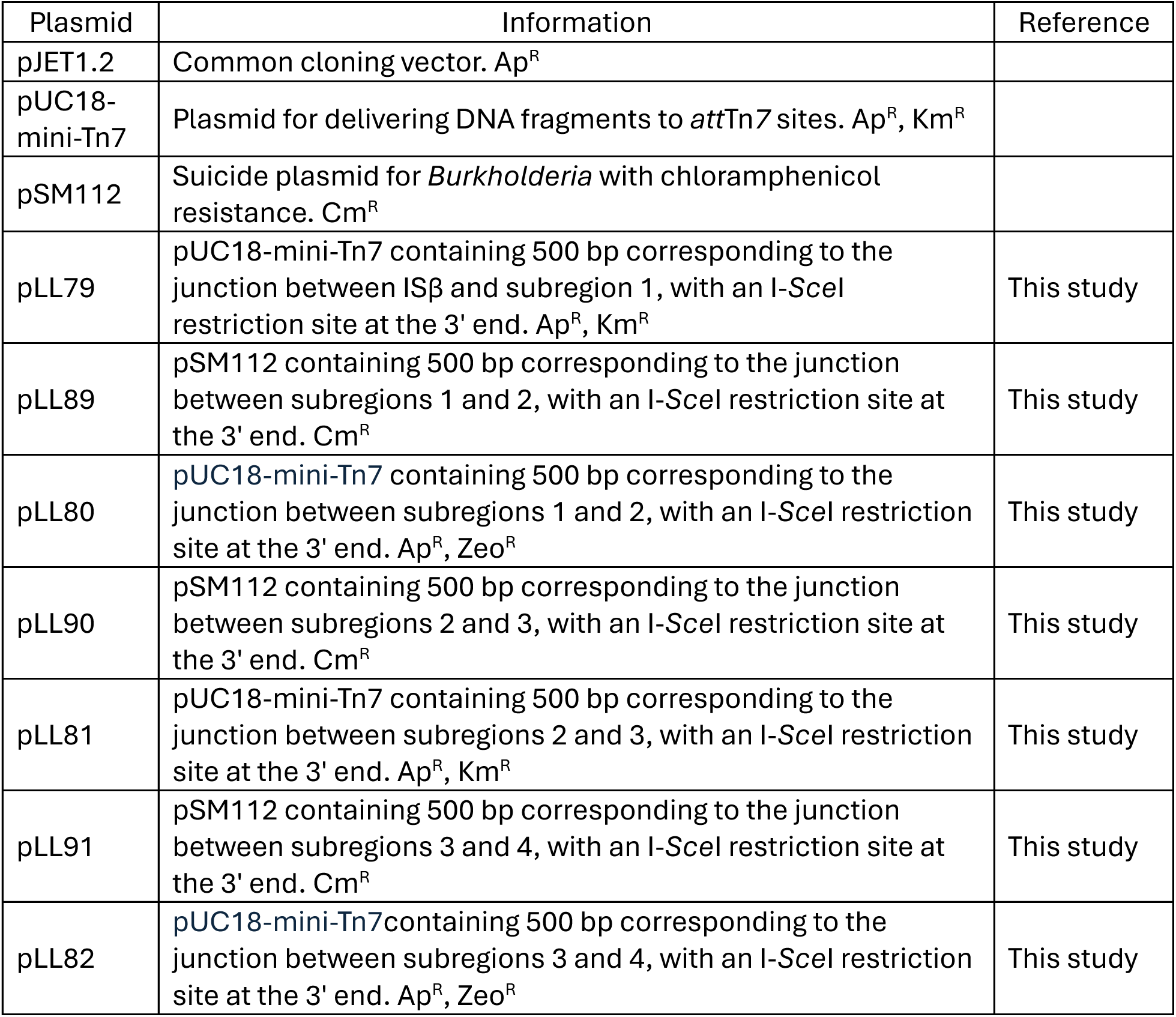
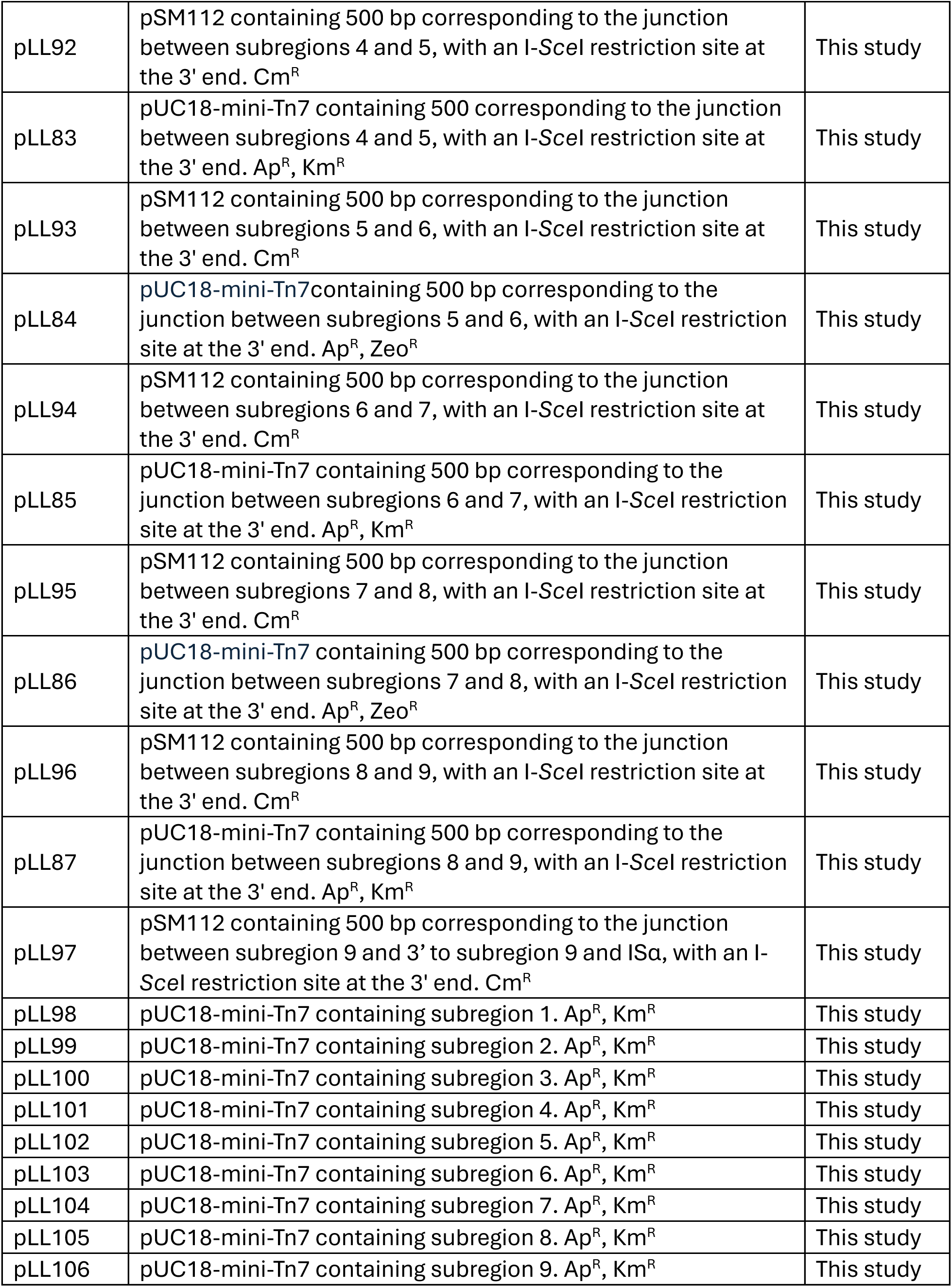
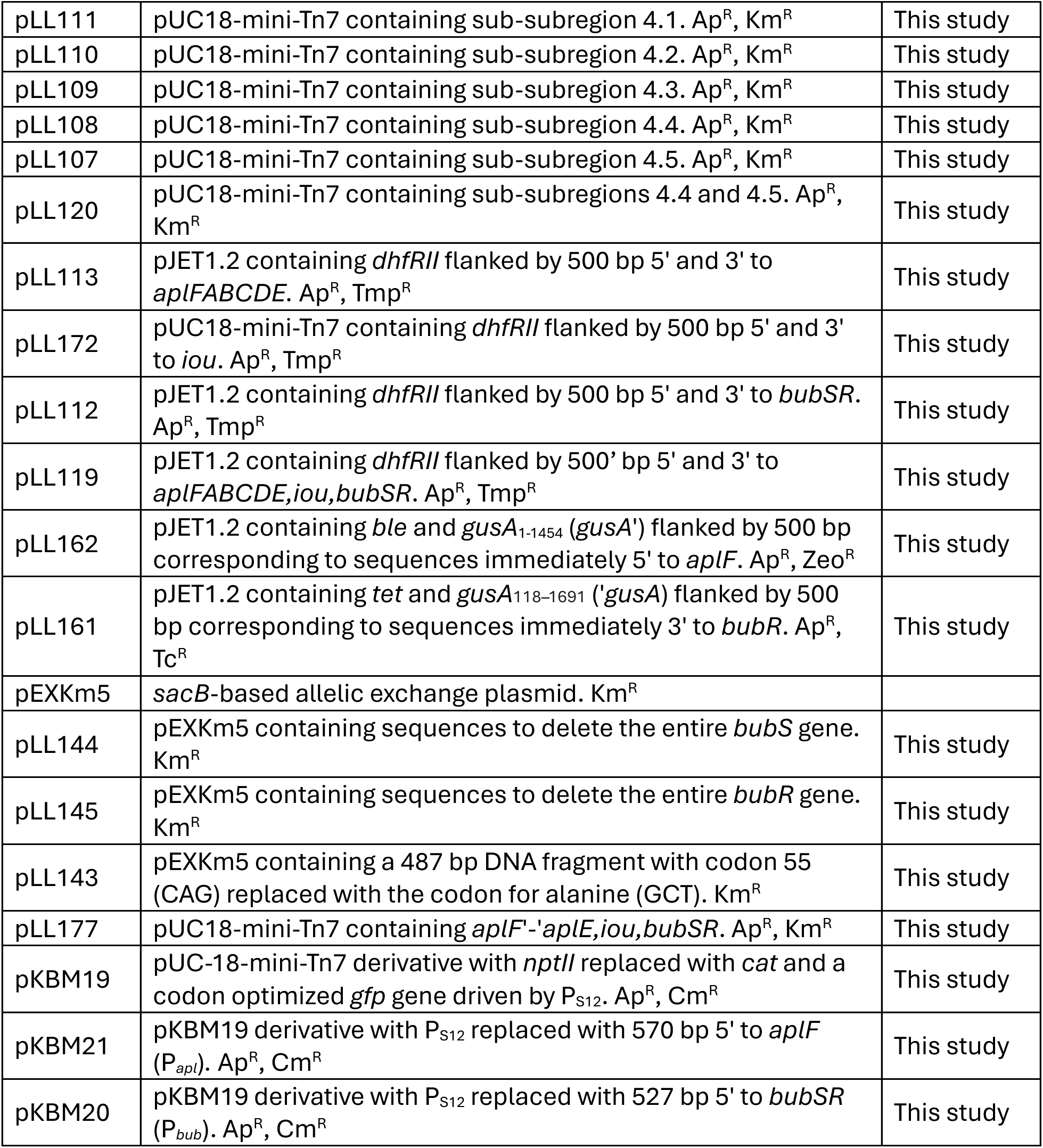

## Strains

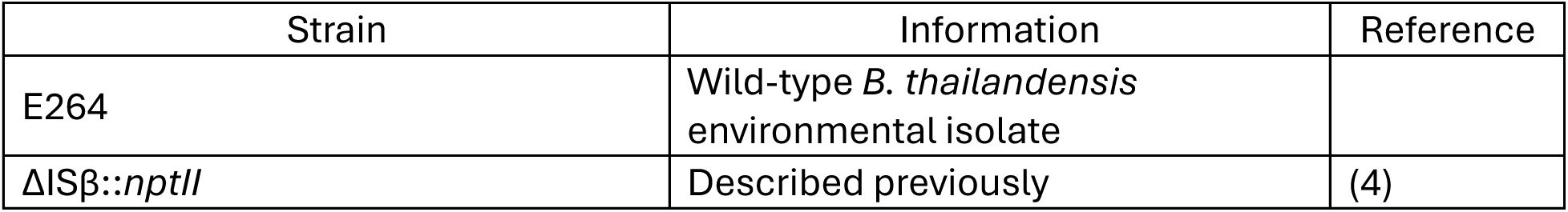
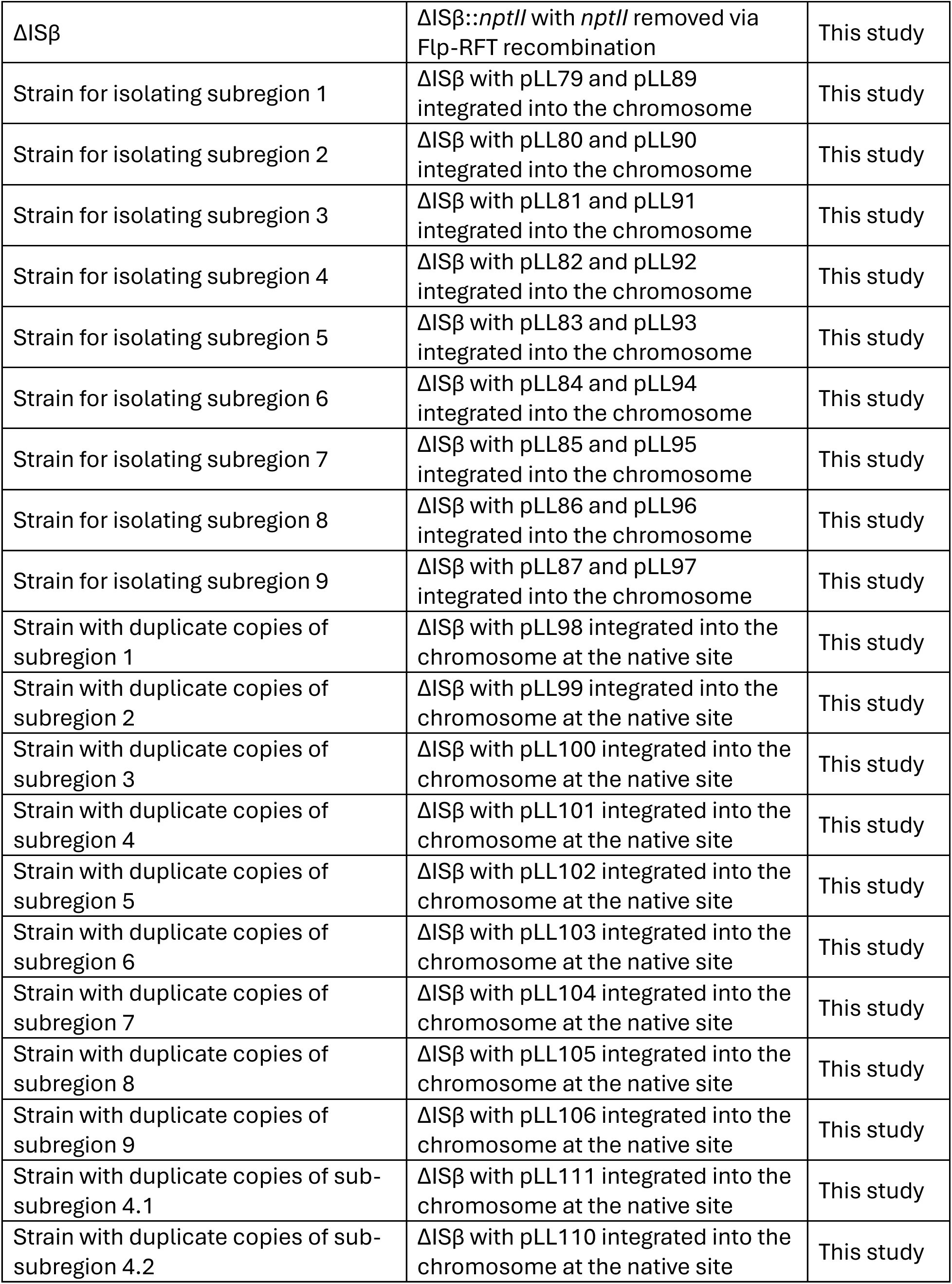
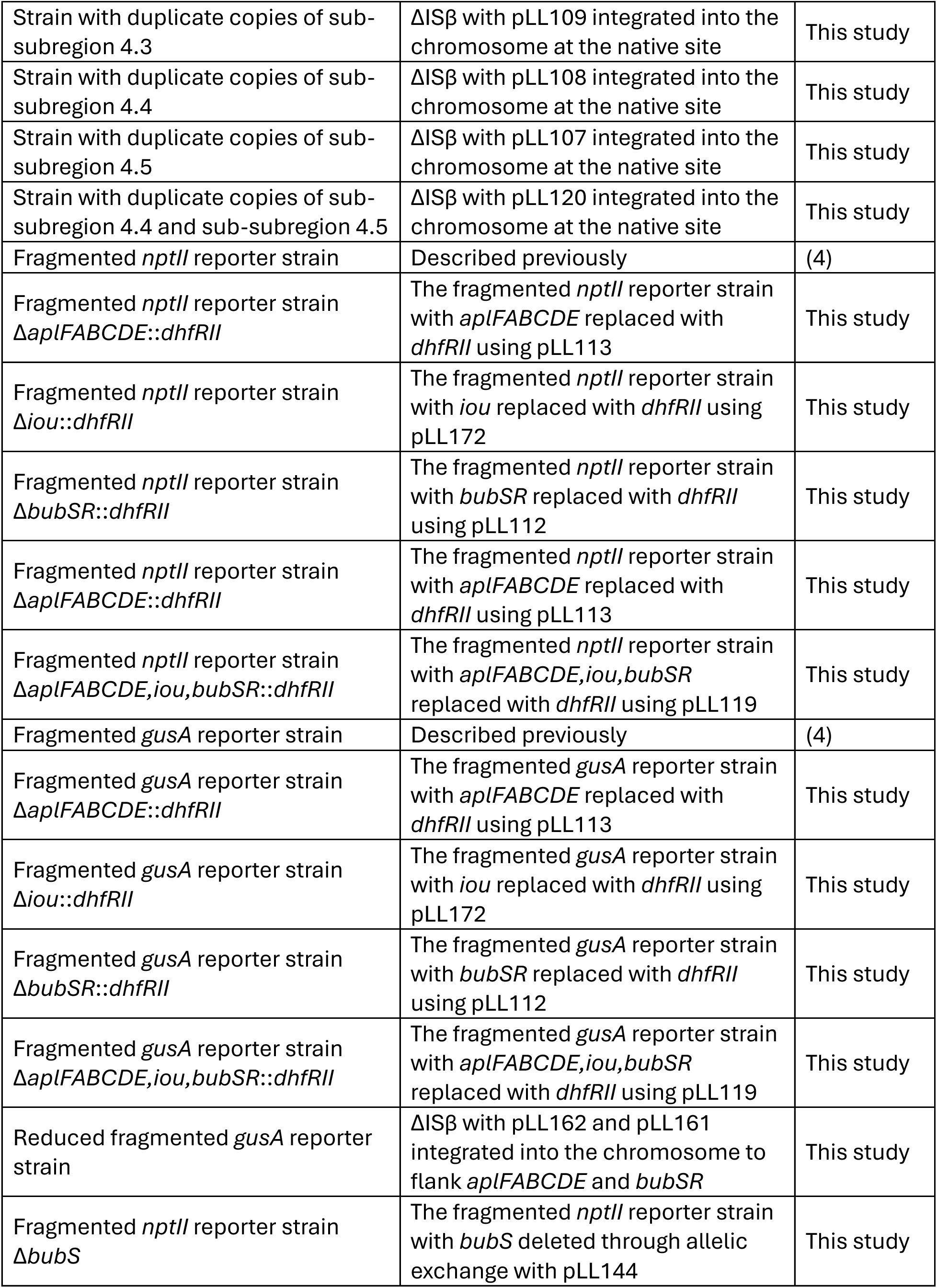
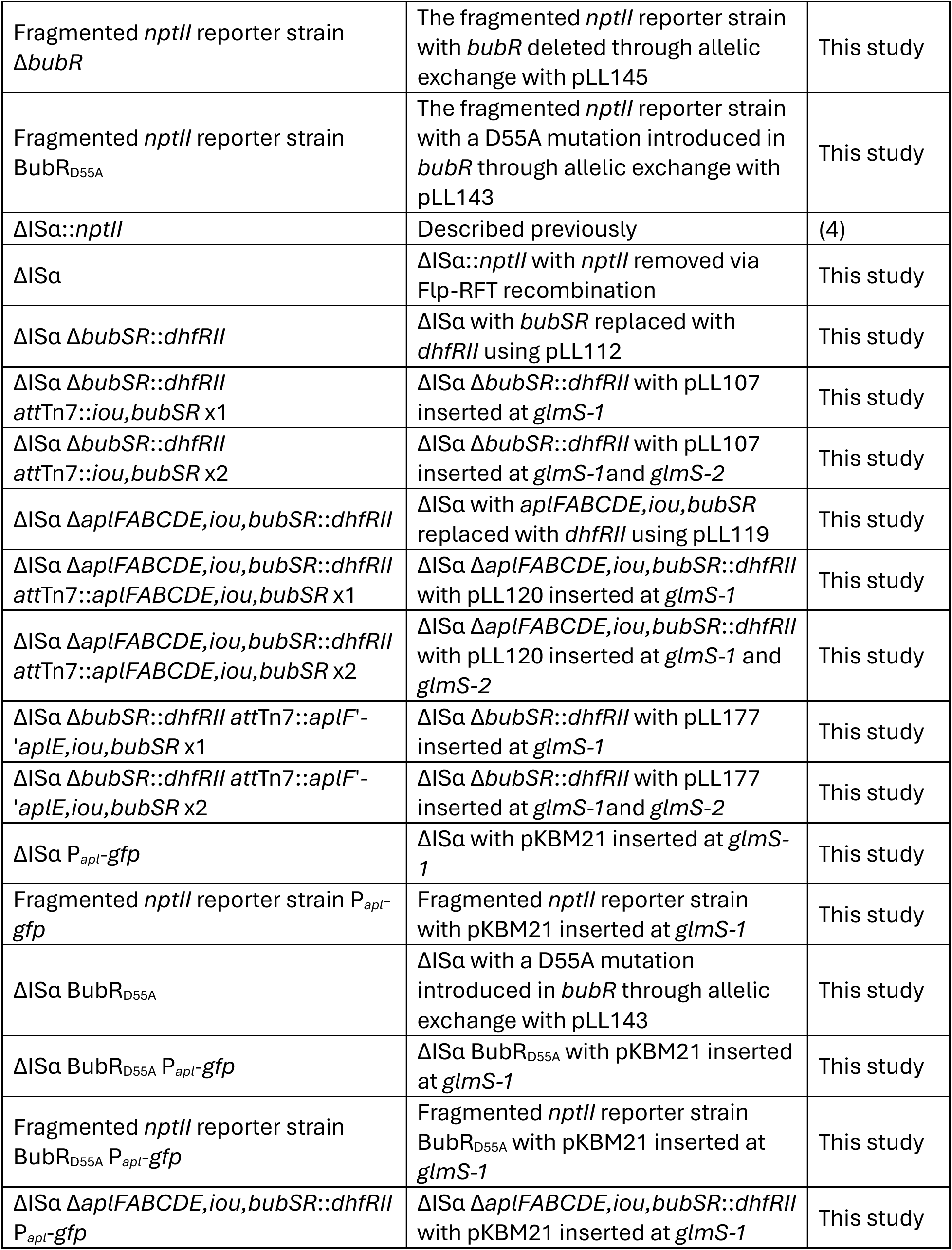
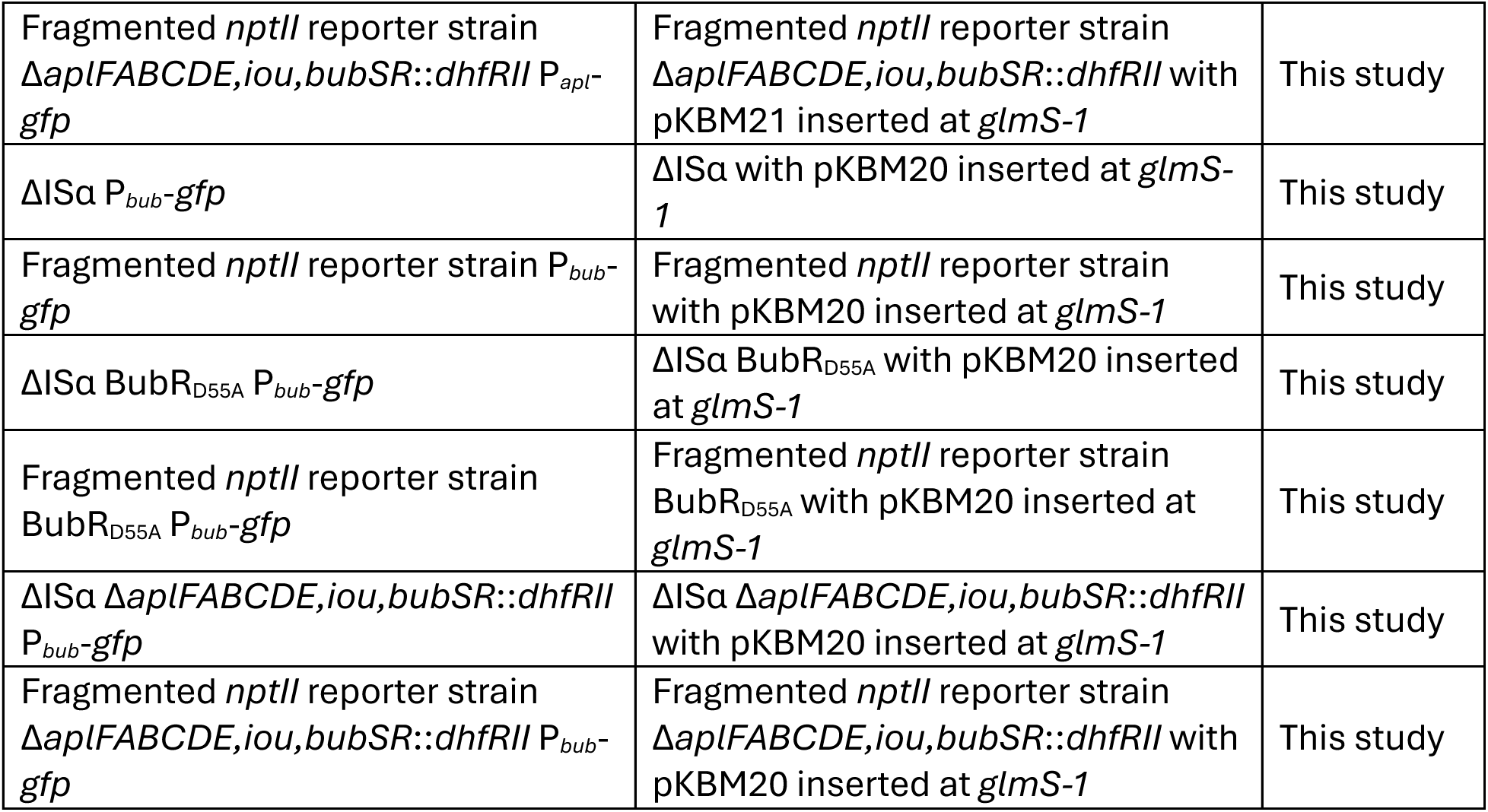

## Acknowledgements

The UNC Flow Cytometry Core Facility (RRID:SCR_019170) is supported in part by P30 CA016086 Cancer Center Core Support Grant to the UNC Lineberger Comprehensive Cancer Center. We thank members of the Cotter Lab, especially Bridgett Rios, for helpful discussions.

---

**Supplemental Figure 1.**
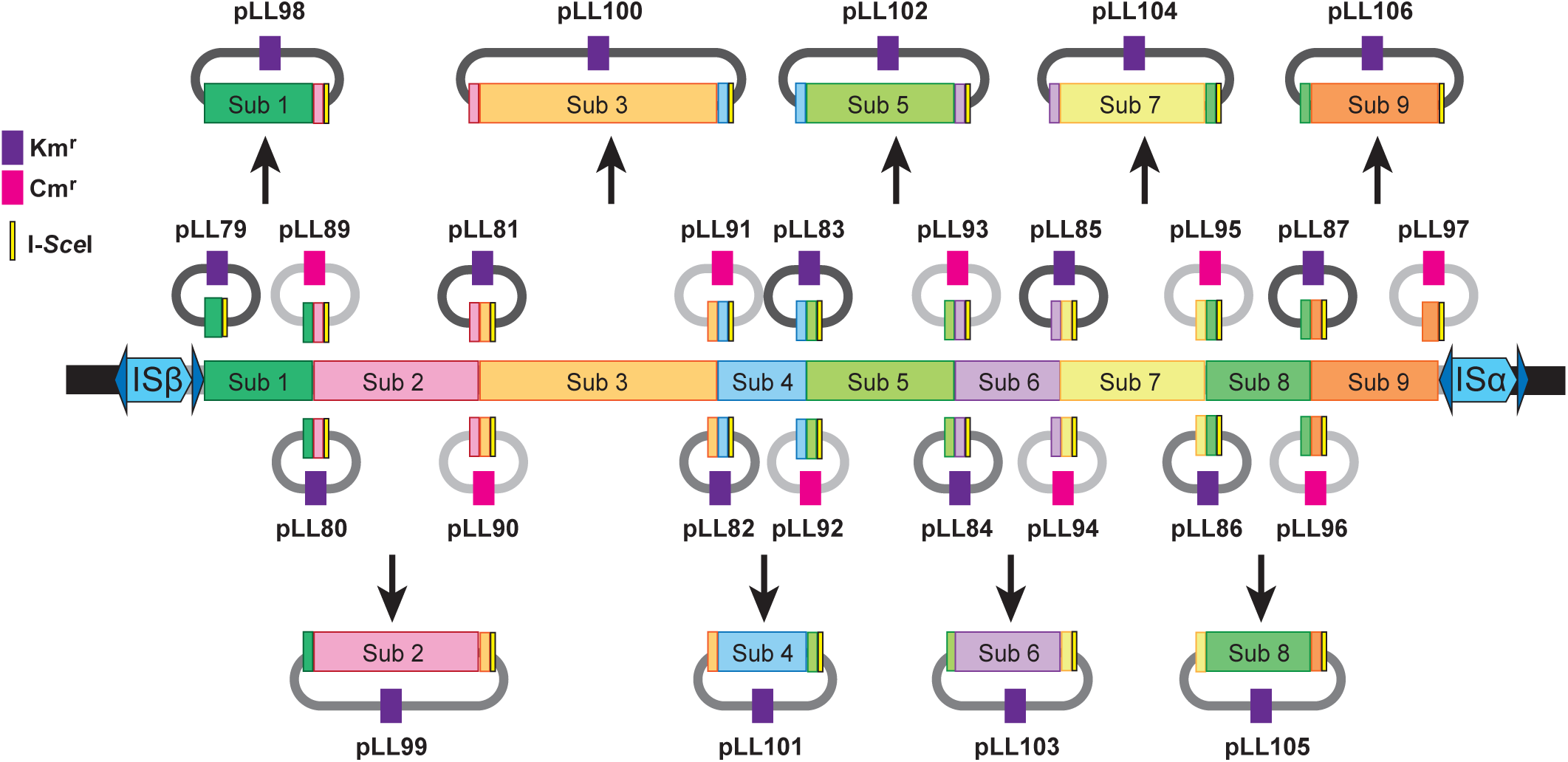
Schematic of modified plasmid rescue strategy used to clone each subregion. The 208.6 kb region was divided into subregions as follows: Subregion 1 contains BTH_I2743 – BTH_I2729 Subregion 2 contains BTH_I2728 – BTH_I2714 Subregion 3 contains BTH_I2713 – BTH_I2687 Subregion 4 contains BTH_I2686 – BTH_I2674 Subregion 5 contains BTH_I2673 – BTH_I2657 Subregion 6 contains BTH_I2656 – BTH_I2641 Subregion 7 contains BTH_I2640 – BTH_I2623 Subregion 8 contains BTH_I2622 – BTH_I2607 Subregion 9 contains BTH_I2606 – BTH_I2587 Each plasmid is a suicide plasmid for *Burkholderia* species and contains a ∼500 bp fragment of DNA corresponding to a junction between subregions, as indicated by the colored boxes, and an I-*Sce*I restriction endonuclease site (yellow box outlined in black), as well as a gene encoding either kanamycin resistance (purple box) or chloramphenicol resistance (dark pink box). Plasmids containing subregions that were generated are shown across the top and bottom.

**Supplemental Figure 2.**
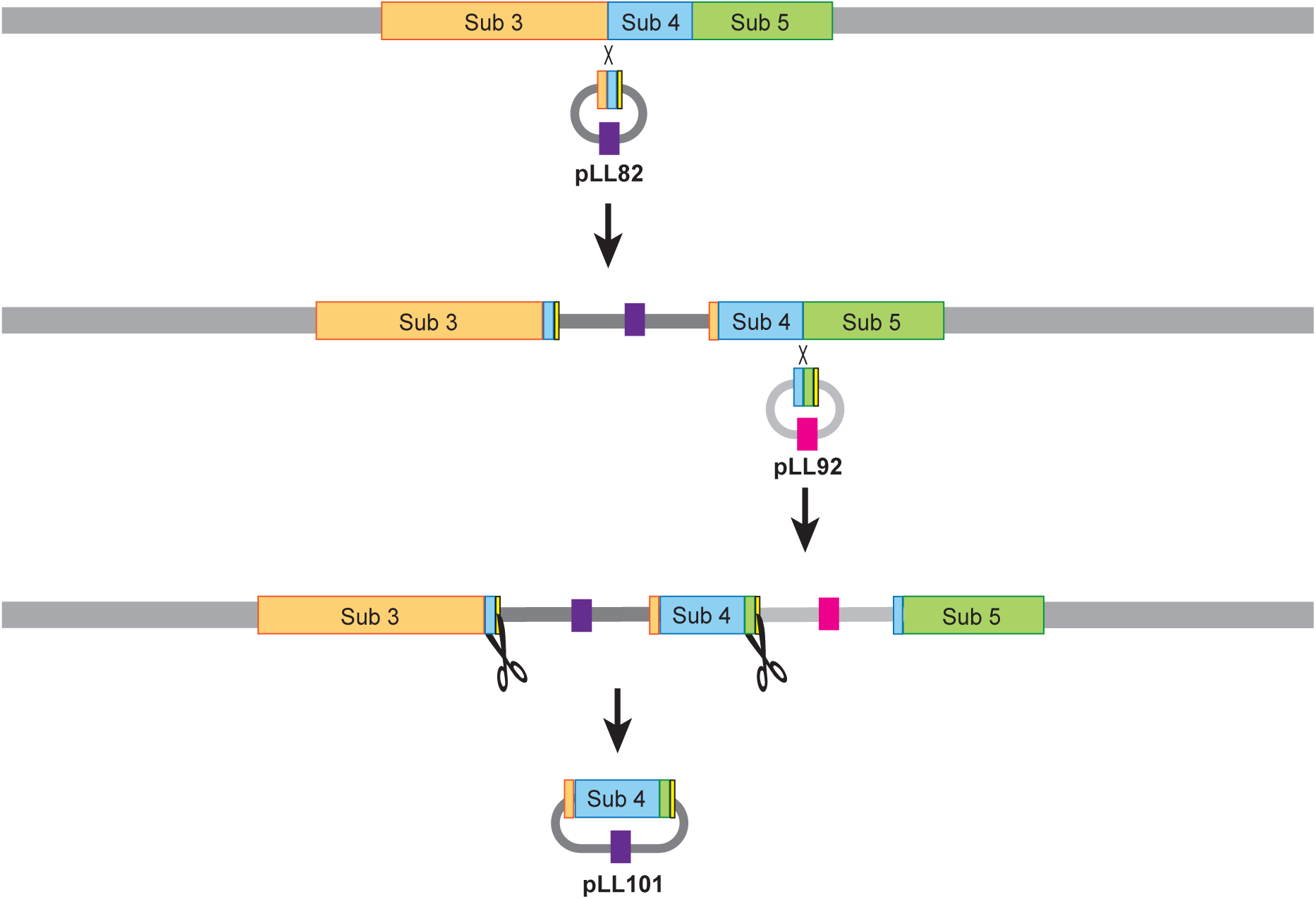
Schematic of the modified plasmid rescue scheme. The plasmid rescue scheme to clone subregion 4 is shown. First, a Kmr cointegrant containing pLL82, which integrates at the junction between subregion 3 and subregion 4 was obtained. Then, a Cmr derivative of that strain containing pLL92 integrated at the junction between subregion 4 and subregion 5 was obtained. Genomic DNA was obtained from the strain containing both cointegrated plasmids, digested with I-*Sce*I and the fragmented DNA was ligated and used to transform E. coli DH5α. *E. coli* colonies were screened for those containing pLL101. All strains and plasmids were confirmed to be as expected by PCR and/or DNA sequence analysis.

**Supplemental Figure 3.**
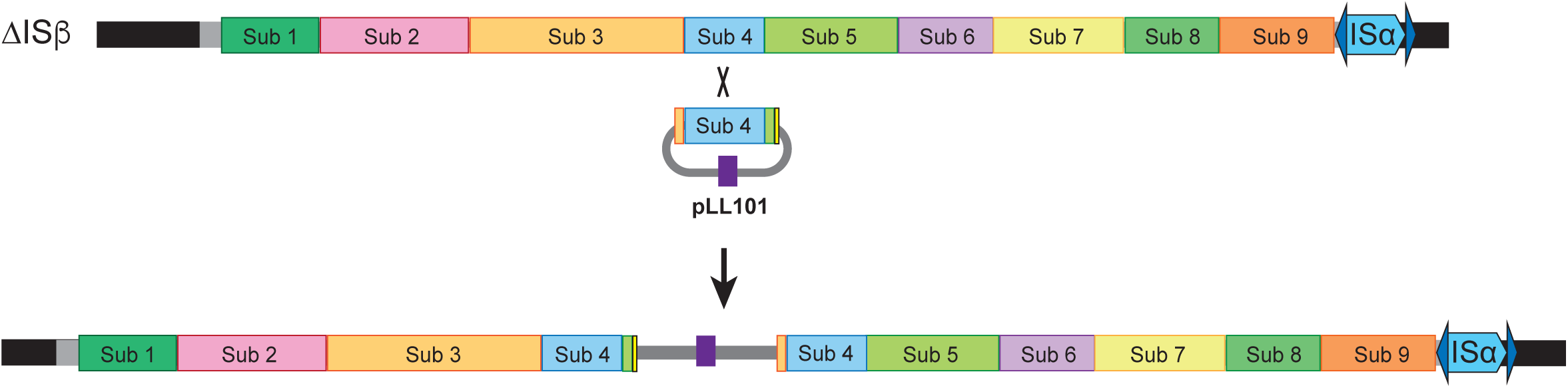
Schematic of constructing strains containing a duplication of only a single subregion. A schematic of constructing a strain containing a duplication of subregion 4 is shown. Plasmid pLL101 was delivered to *Bt*E264 ΔISβ, which cannot duplicate the 208.6 kb region, by conjugation and Kmr cointegrants were obtained. Integration of pLL101 into subregion 4 was confirmed by PCR.

